# Phosphodiesterase 3 inhibitors boost bone outgrowth

**DOI:** 10.1101/2024.07.01.601482

**Authors:** Takaaki Kawabe, Atsuhiko Ichimura, Tomoki Yasue, Jianhong Li, Ga Eun Kim, Haruki Ishikawa, Yohei Uda, Hiromu Ito, Miyuki Nishi, Hiroshi Takeshima

**Author notes:** Corresponding Authors: Atsuhiko Ichimura, Laborarory of Integrative Physiology, Collage of Pharmaceutical Sciences, Ritsumeikan University, Shiga 525-8577, Japan.; Hiroshi Takeshima, Graduate School of Pharmaceutical Sciences, Kyoto University, Kyoto 606-8501, Japan. These authors equally contributed to this study (Takaaki Kawabe, Atsuhiko Ichimura).

## Abstract

C-type natriuretic peptide (CNP) stimulates skeletal growth by acting on the growth plates of long bones, and a CNP variant is clinically used for achondroplasia treatment. We previously reported that CNP stimulates the autonomic Ca^2+^ influx mediated by TRPM7 channels in growth plate chondrocytes to facilitate extracellular matrix synthesis for bone growth. Here, we report that phosphodiesterase 3 (PDE3) inhibitors facilitate the Ca^2+^ influx and promote bone outgrowth. The representative PDE3 inhibitor cilostazol elevated cGMP levels and activated cell-surface K^+^ channels probably due to protein kinase G-mediated phosphorylation in growth-plate chondrocytes. The resulting hyperpolarization likely facilitated TRPM7-mediated Ca^2+^ influx by increasing the Ca^2+^-driving force. Moreover, cilostazol simulated the elongation of cultured bones and enlarged the body size of juvenile mice. Several PDE3 inhibitors have been used for clinical treatment of thrombosis, heart failure and asthma, while our observations suggest that the repositioning of PDE3 inhibitors would provide novel medications for syndromes involving short stature.

## INTRODUCTION

Skeletal long bones develop through endochondral ossification, during which primitive chondrocytes initially form growth plates at both ends of bone rudiments (Mackie *et al*, 2008). The growth plates then form three distinct zones, each of which contains chondrocytes that are homogenous in cell shape and maturation stage: “round chondrocytes” differentiated from chondroblasts at most ends, “columnar chondrocytes” characterized by a unique flat shape, and nonproliferating “hypertrophic chondrocytes” undergoing apoptosis at the bone rudiment center (Berendsen & Olsen, 2015). Bone growth mainly depends on the vital proliferation and characteristic extracellular matrix (ECM) synthesis of proliferating round and columnar chondrocytes, while genetic and environmental factors naturally affect cellular functions. In the hypertrophic chondrocyte zone, the region containing apoptotic bodies is gradually replaced by trabecular bones through the actions of osteoclasts and osteoblasts during juvenile development (Mackie *et al*., 2008).

The peptide hormone C-type natriuretic peptide (CNP) acts on proliferating chondrocytes expressing the CNP-specific receptor guanylate cyclase NPR2 in an autocrine/paracrine manner and promotes long bone growth(Kake *et al*, 2009; Nakao *et al*, 2009; Suga *et al*, 1992; Teixeira *et al*, 2008; Yasoda *et al*, 2004). Accordingly, loss- and gain-of-function mutations in the *NPR2* gene were reported to cause acromesomelic dysplasia and skeletal overgrowth disorder, respectively (Bartels *et al*, 2004; Khan *et al*, 2016; Robinson *et al*, 2013). Furthermore, the CNP variant peptide vosoritide is used for the clinical treatment of achondroplasia, which is characterized by short limb bones and is mainly caused by gain-of-function mutations in the fibroblast growth factor receptor 3 (*FGFR3*) gene (Legeai-Mallet, 2016; Savarirayan *et al*, 2020).

In the transient receptor potential channel superfamily, TRPM7 (melastatin subfamily member 7) forms a cation-permeable channel and participates in divergent cellular processes including cell growth and adhesion in various cell types (Fleig & Chubanov, 2014). We previously reported that TRPM7 channels generate autonomic Ca^2+^ influx in proliferating growth plate chondrocytes, leading to the activation of Ca^2+^/calmodulin-dependent protein kinase II (CaMKII), which stimulates ECM synthesis and thus facilitates bone outgrowth (Qian *et al*, 2019). In addition, we revealed that the following CNP signaling axis is essential for facilitating bone outgrowth (Miyazaki *et al*, 2022). CNP-induced NPR2 activation elevates the cellular cGMP content and activates cGMP-dependent protein kinase (PKG), leading to the phosphorylation and activation of big-conductance Ca^2+^-dependent potassium (BK) channels in growth plate chondrocytes. The resulting BK channel activation induces hyperpolarization to facilitate TRPM7-mediated Ca^2+^ entry by enhancing the Ca^2+^ driving force and thus stimulating CaMKII. The defined cGMP-driven signaling cascade, the PKG-BK channel-TRPM7 channel-CaMKII axis, seems to pinpoint target proteins for developing new therapeutic treatments for skeletal growth disorders (Fig. EV1). In this context, we attempted to increase the steady-state cGMP concentration using phosphodiesterase (PDE) inhibitors to promote CNP signaling. In this report, we demonstrate that a class of PDE inhibitors activates the CNP signaling to promote spontaneous Ca^2+^ influx in growth plate chondrocytes and thus stimulates long bone growth.

## RESULTS

### PDE subtypes in growth plate chondrocytes

The aim of this study was to determine whether chemical PDE inhibitors can artificially stimulate bone growth by facilitating the CNP signaling in growth plate chondrocytes. Among the PDE family subtypes, fifteen are presumed to physiologically contribute to cGMP inactivation in mammalian cell types (Heckman *et al*, 2018; Omori & Kotera, 2007; Stratakis *et al*, 2014). Initially, we attempted to identify the PDE candidates that control the steady-state cGMP level in proliferating growth plate chondrocytes by microarray and RT-PCR analyses using RNA preparations from mouse embryonic femoral bones. Our gene expression data suggested that the cGMP-hydrolyzing subtype genes *Pde1b*, *Pde2a*, *Pde3b, Pde5a*, *Pde6g*, *Pde9a* and *Pde10a* were moderately expressed in round chondrocytes (Fig. 1). PDE1B is Ca^2+^/calmodulin-dependently activated to hydrolyze both cAMP and cGMP (Samidurai *et al*, 2021), while PDE2A is activated by cGMP to balance cellular cAMP and cGMP levels (Sadek *et al*, 2020). Based on their biological features, PDE1B and PDE2A are unlikely to contribute to establishing the basal cGMP concentration under resting conditions. *Pde5a-*knockout mice exhibit small body sizes (Data ref: International Mouse Phenotyping Consortium MGI: 2651499, 2011), and *Pde9a*-knockout mice have normal body sizes (Ceddia *et al*, 2021), suggesting that inhibitors of PDE5A and PDE9A are unlikely to stimulate bone growth. *Pde6g*-knockout mice develop retinitis pigmentosa blindness but have normal body sizes (Tsang *et al*, 1996). Human *PDE6G* and *PDE10A* mutations are responsible for blindness and parkinsonism, respectively, while no skeletal abnormalities have been found in either the disease cases (Dvir *et al*, 2010; Siuciak *et al*, 2008). Therefore, PDE6G and PDE10A may not contribute to major bone growth signaling pathways. The processes of elimination highlighted PDE3B in the proliferating chondrocytes, and it is also notable that *Pde3b*-knockout mice exhibit the enlargement of the typical long bone tibia as a skeletal phenotype (Data ref: International Mouse Phenotyping Consortium MGI:1333863, 2011). We thus focused on the pharmacological effects of clinically used PDE3 inhibitors on long bone development.

**Fig. 1.**
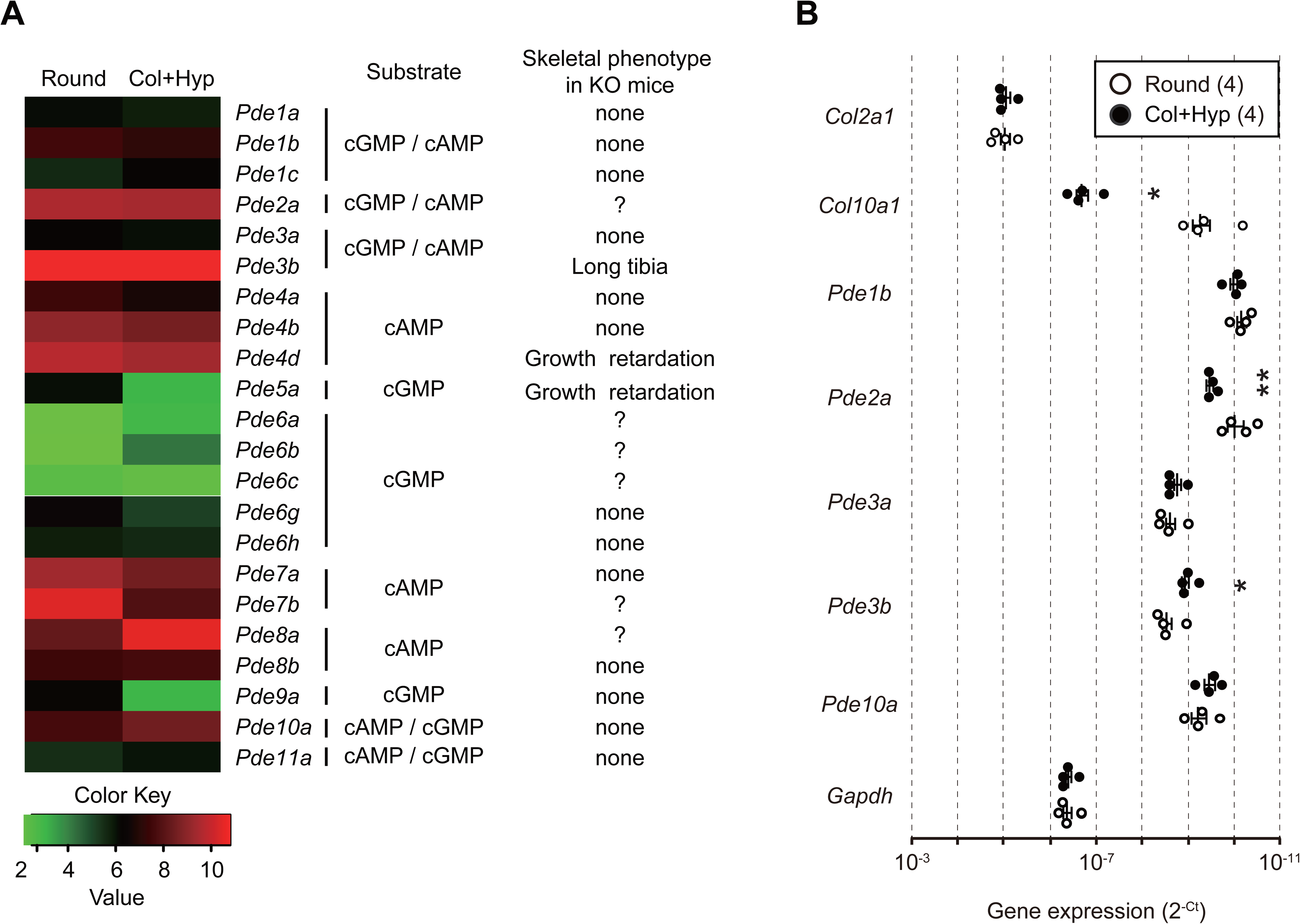
Gene expression analysis in growth plate chondrocytes. **(A)** Microarray heatmap representing the expression levels of PDE subtype genes in mouse growth plates. Growth plate sections enriched in either round chondrocytes (Round) or columnar and hypertrophic chondrocytes (Col+Hyp) were dissected from E17.5 femoral bones and subjected to total RNA preparation for microarray analysis. The resulting gene expression values are represented according to the color scale. The substrate specificity and skeletal phenotype of each PDE subtype observed in knockout mice are also listed. **(B)** RT-PCR analysis of PDE subtype genes in growth plate sections in either round chondrocytes or columnar and hypertrophic chondrocytes. The primer sets used are listed in Table EV2 and the cycle threshold (Ct) was determined for each RT-PCR amplification. The data are presented as the mean ± SEM., and the numbers of mice examined are shown in parentheses. Statistical differences between the growth plate sections are indicated by asterisks (* *p*<0.05, ** *p* < 0.01 according to t test).

### PDE3 inhibitors facilitate cultured bone growth

*Ex vivo* metatarsal culture has been exploited as a useful method for analyzing endochondral bone growth (Staines *et al*, 2019). We examined the effects of PDE inhibitors supplemented to culture medium on the longitudinal elongation of metatarsal bones prepared from embryonic mice. The representative PDE3 inhibitor cilostazol significantly stimulated the bone outgrowth during 4 days of culture (Fig. 2A); the bone length increased by 120.7 ± 0.9% and 127.6 ± 1.2% in the control medium and 10 μM cilostazol-containing medium, respectively. The other PDE3 inhibitors milrinone, anagrelide and olprinone also promoted cultured bone elongation, while the PDE2 inhibitor BAY-60-7550 and the PDE10 inhibitor MP10 had no effects (Fig. EV2 and EV3A). Mutant mice carrying the chondrocyte-specific *Fgfr3* transgene have been used as an animal model of achondroplasia (Wang *et al*, 1999), and cilostazol supplementation was also effective in the mutant bones from the transgenic mice (Fig. EV4A). Therefore, PDE3 inhibitors seemed to commonly stimulate bone outgrowth in *ex vivo* culture.

**Fig. 2.**
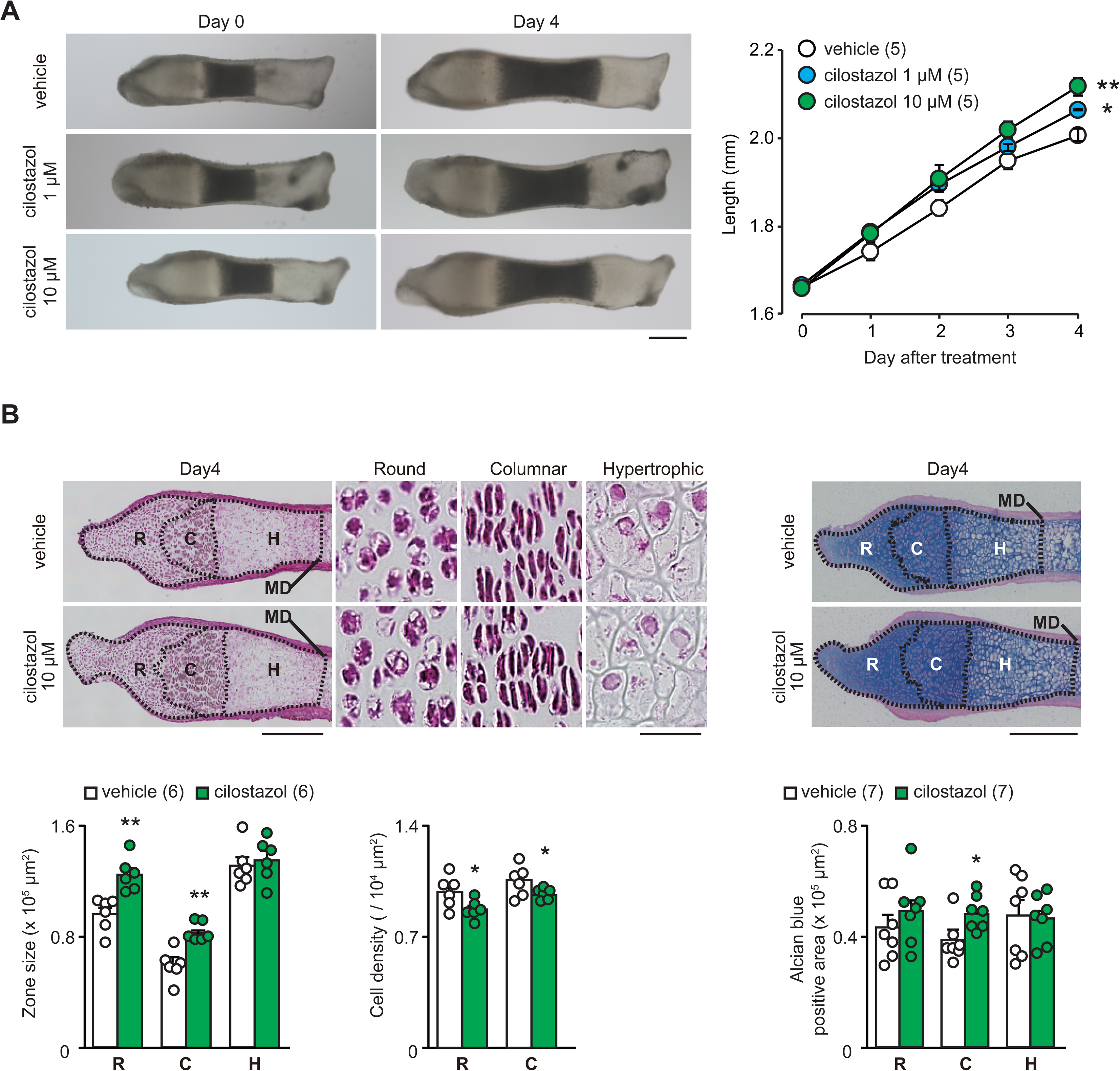
Cilostazol promotes cultured bone growth. **(A)** Cilostazol-promoted outgrowth of cultured metatarsal bones. Representative images of the metatarsals before and after culture for 4 days in the presence or absence of cilostazol (left panels; scale bar, 0.3 mm). Longitudinal growth of the metatarsals was quantified over the culture period (right graph). The numbers of mice examined are shown in parentheses. The bone lengths at the endpoint were analyzed and statistical differences from the vehicle-treated group are marked with asterisks (\**p*<0.05 and \*\**p*<0.01 according to two-way ANOVA and Dunnett’s test). **(B)** Histological analysis of metatarsal bones cultured in the presence or absence of cilostazol. The mid-longitudinal sections of cultured metatarsal bones were stained with hematoxylin and eosin (upper left panels; scale bar, 0.3 mm), and their high-magnification images of the round (R), columnar (C), and hypertrophic (H) chondrocyte zones are also shown (scale bar, 30 μm). The metatarsal sections were likewise stained with alcian blue (upper right panels; scale bar, 0.3 mm). MD indicates the mid-diaphysis. In the round, columnar and hypertrophic chondrocyte zones, the zonal size, cell density and alcian blue-positive area were examined and are summarized (lower graphs). The data are presented as the mean ± SEM., and the numbers of mice examined are shown in parentheses. Statistical differences between the vehicle- and cilostazol-treated groups are marked with asterisks (\**p* < 0.05 and \*\**p* < 0.01 according to t test).

Longitudinal sections were prepared from the cultured metatarsals and subjected to histological analysis of growth plates. In the cilostazol-treated bones, round and columnar chondrocyte zones were preferentially expanded (Fig. 2B). Although the cell densities were slightly decreased, alcian blue-positive ECM regions tended to be enlarged in the extended proliferating chondrocyte zones. The milrinone-treated bones also exhibited histological features essentially the same as those of the cilostazol-treated bones (Fig. EV3B). Therefore, PDE3 inhibitors likely stimulated ECM production in proliferating chondrocytes to facilitate growth plate outgrowth toward bone elongation. Furthermore, the histological alterations induced by PDE3 inhibitors were essentially similar to those of CNP-treated cultured bones that we previously observed (Miyazaki *et al*., 2022).

### Cilostazol stimulates mouse body growth

To examine the *in vivo* effects of PDE3 inhibitors on body growth, juvenile mice immediately after delactation (3-weeks old) were daily injected with cilostazol at 10 mg/kg (Fig. 2A). During the 4-week test period, body size (naso-anal length) increased by 93.6 ± 0.3 mm in the vehicle-treated control mice and increased by 95.3 ± 0.5 mm in the cilostazol-treated mice. Therefore, cilostazol application potentiated body growth in young mice. After the test period, tibial sections were prepared from the mice and subjected to histological analysis (Fig. 3B). As with the cultured metatarsal bones, the alcian-blue-positive ECM regions were expanded in the enlarged growth plates prepared from the cilostazol-treated mice (Fig. 3C). Although cilostazol frequently causes hypotension and tachycardia as its side effects, the cilostazol treatment seemed to exert no influence on the major cardiovascular or hematological parameters during the test period (Fig. EV5). Based on the observations that PDE3 inhibitors stimulated cultured bone growth and juvenile mouse growth, we next focused on the pharmacological effects of cilostazol on growth plate chondrocytes.

**Fig. 3.**
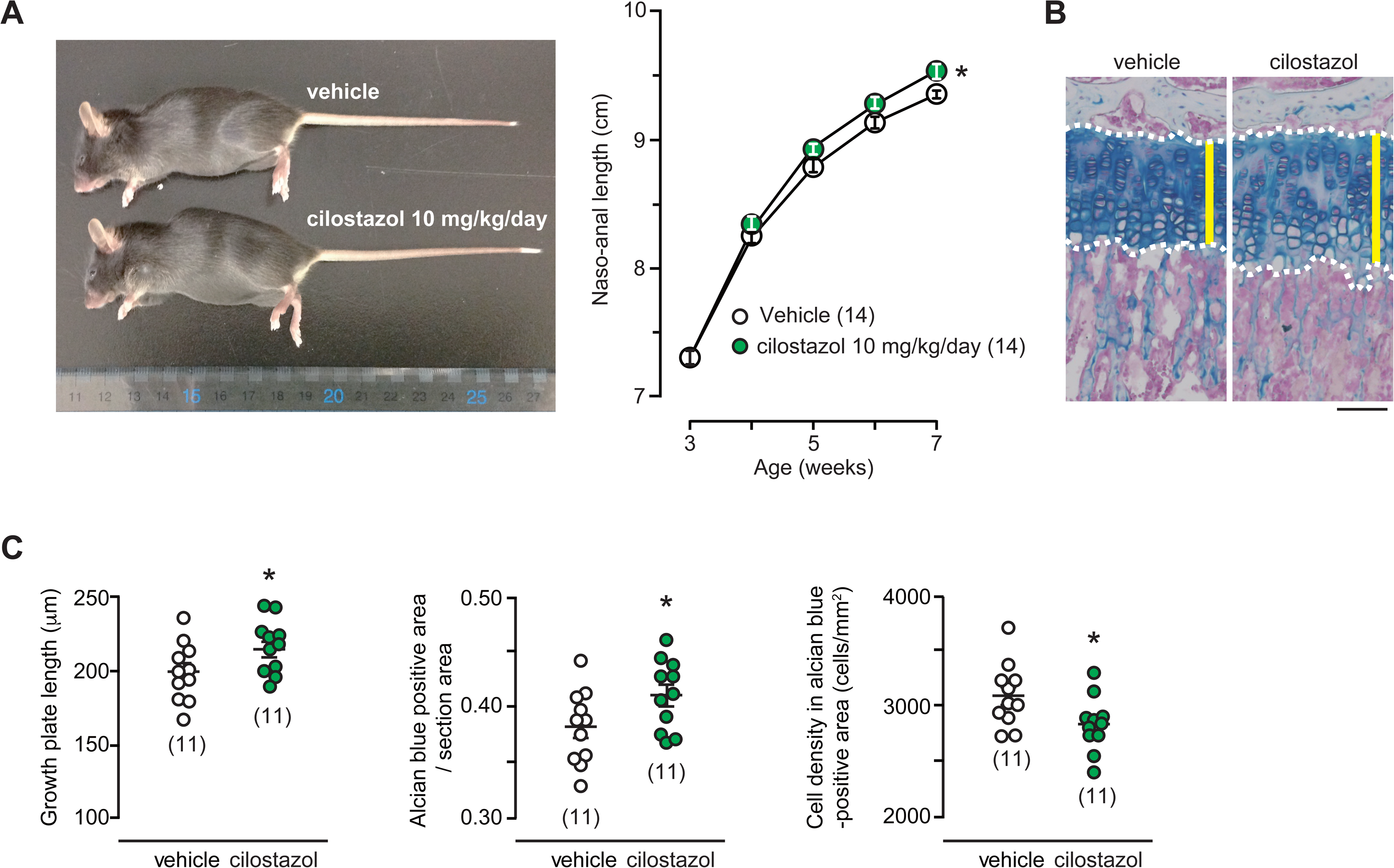
Cilostazol stimulates body growth in juvenile mice. **(A)** Increased body length by daily intraperitoneal injection of cilostazol. Representative images of the mice treated with vehicle or cilostazol for 4 weeks (left panel). Transitions in naso-anal length were analyzed over the test period and the data are summarized (right graph). The numbers of mice examined are shown in parentheses, and statistical differences at the endpoint are marked with asterisks (\**p* < 0.05 according to two-way ANOVA and Sidak’s test). **(B)** Representative images of alcian blue-stained tibial growth plates from the vehicle- and cilostazol-treated mice. The mice were drug-treated for 2 weeks (3 to 5-week-old) and then subjected to the histological analysis. Yellow bars indicate alcian blue-positive growth plate zone. Scale bar, 0.1 mm. **(C)** Tibial growth plate length, alcian blue-positive area per section area, and cell density in the alcian blue-positive area are summarized. The data are presented as the mean ± SEM., and the numbers of mice examined are shown in parentheses. Significant differences between the groups are indicated by asterisks (\**p* < 0.05 according to t test).

### Cilostazol elevates cGMP levels in growth plate chondrocytes

PDE3B hydrolyzes both cAMP and cGMP, but its cAMP hydrolytic activity is attenuated by elevated cGMP content (Omori & Kotera, 2007). To determine the fundamental role of PDE3B in growth plate chondrocytes, we isolated femoral bones from embryonic mice and prepared the epiphyseal slices enriched in proliferating chondrocytes for the immunochemical quantification of cAMP and cGMP levels. In response to cilostazol treatments to the slice specimens (3 μM for 30 min), the cGMP content increased ∼1.7-fold, while the cAMP content remained unchanged (Fig. 4A). Similar effects on the cyclic nucleotide content were also observed in the milrinone treatment group (Fig. EV6A). Therefore, PDE3 inhibitors seemed to selectively elevate cGMP concentration in growth plate chondrocytes.

**Fig. 4.**
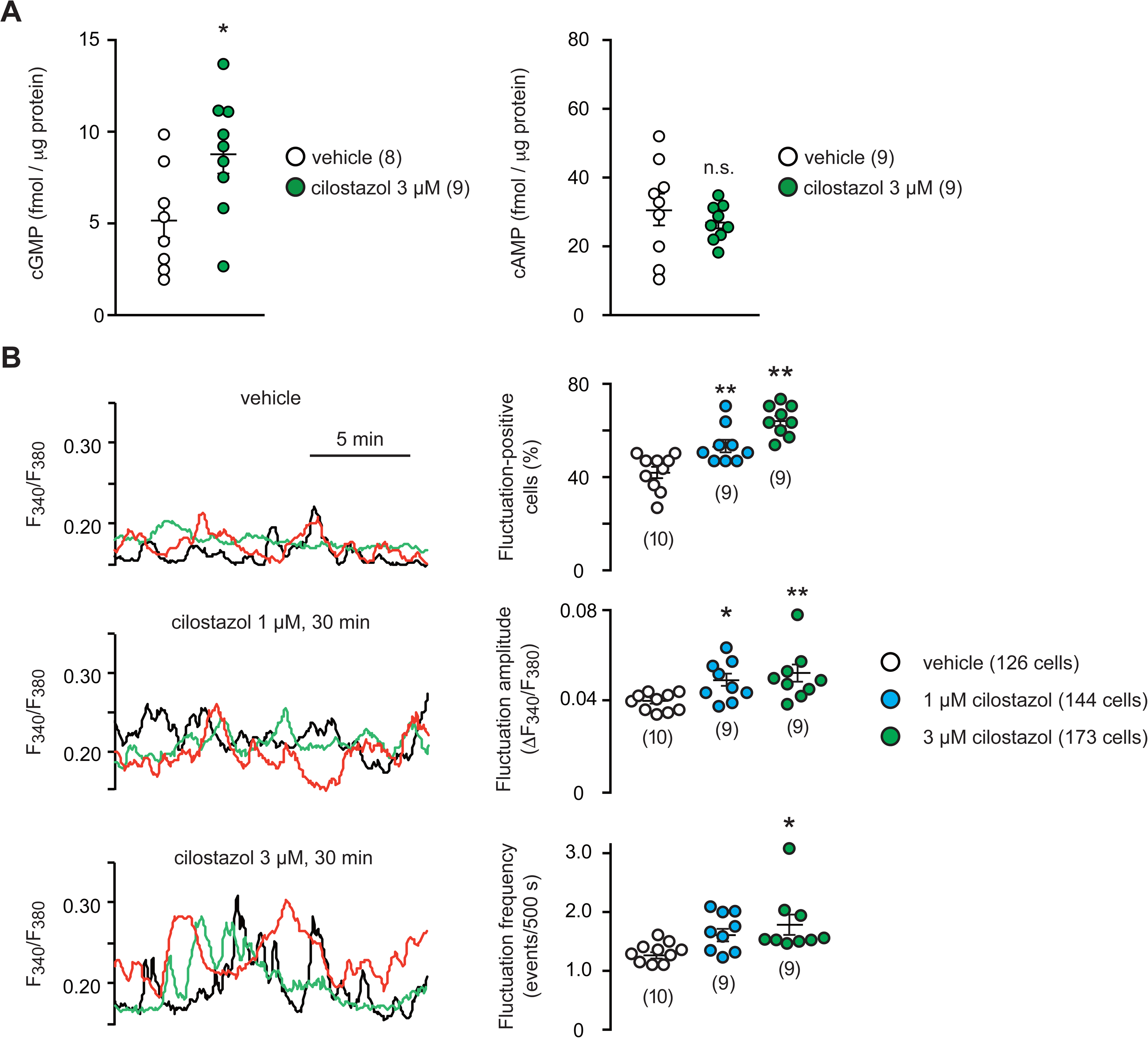
Cilostazol facilitates Ca^2+^ fluctuations in growth plate chondrocytes. **(A)** Effects of cilostazol on cGMP and cAMP levels in growth plate chondrocytes. Femoral bone slices enriched in growth plate chondrocytes were treated with or without cilostazol for 30 min and then subjected to cyclic nucleotide quantification. The data are presented as the mean ± SEM., and the numbers of mice examined are shown in parentheses. A significant difference between the groups is indicated by an asterisk (\**p* < 0.05 according to t test; n.s., not significant). **(B)** Effects of cilostazol on Ca^2+^ fluctuations in growth plate chondrocytes. Fura-2 imaging of round chondrocytes in femoral bone slices pretreated with or without cilostazol for 30 min. Representative recording traces from three cells are shown for each pretreatment group (left panels). The effects of cilostazol on spontaneous Ca^2+^ fluctuations are summarized (right graphs); the fluctuation-positive cell ratio, fluctuation amplitude and fluctuation frequency were statistically analyzed. The numbers of mice and cells examined are shown in the bar graphs and keys, respectively. Statistical differences from the vehicle-pretreated group are marked with asterisks (\**p* < 0.05 and ** *p* < 0.01 according to one-way ANOVA and Dunnett’s test).

### Cilostazol stimulates Ca^2+^ entry in growth plate chondrocytes

In proliferating growth plate chondrocytes, TRPM7 channels spontaneously and intermittently open to generate intracellular Ca^2+^ fluctuations, and the TRPM7-mediated Ca^2+^ influx is facilitated by BK channel-induced hyperpolarization (Qian *et al*., 2019). We next analyzed the effects of cilostazol pretreatment (1 or 3 μM for 30 min) on the Ca^2+^ fluctuations in round chondrocytes. In Fura-2 Ca^2+^ imaging of femoral bone slices, we observed that the cilostazol treatments remarkably facilitated the Ca^2+^ fluctuations; the fluctuation-positive cell rate, fluctuation amplitude and frequency were dose-dependently increased in the cilostazol-treated chondrocytes (Fig. 4B). Furthermore, milrinone exerted the effects on the Ca^2+^ fluctuations essentially similar to those of cilostazol (Fig. EV6B). Therefore, PDE3 inhibitors seemed to commonly facilitate the Ca^2+^ fluctuations in proliferating chondrocytes.

The PKG inhibitor KT5823 and the BK channel inhibitor paxilline obviously attenuated the cilostazol-facilitated Ca^2+^ fluctuations (Fig. 5A and B), suggesting the involvement of the PKG and BK channels in the cilostazol-evoked signaling pathway. The TRPM7 inhibitor FTY720 almost abolished the facilitated Ca^2+^ fluctuations (Fig. 5C), indicating that TRPM7-mediated Ca^2+^ entry was facilitated by cilostazol. Based on these observations, together with the cyclic nucleotide quantification data, we could reasonably propose that PDE3 inhibitors increased the cGMP content and stimulated PKG to induce BK channel phosphorylation, leading to the facilitation of TRPM7-mediated Ca^2+^ influx by inducing hyperpolarization in proliferating chondrocytes.

**Fig. 5.**
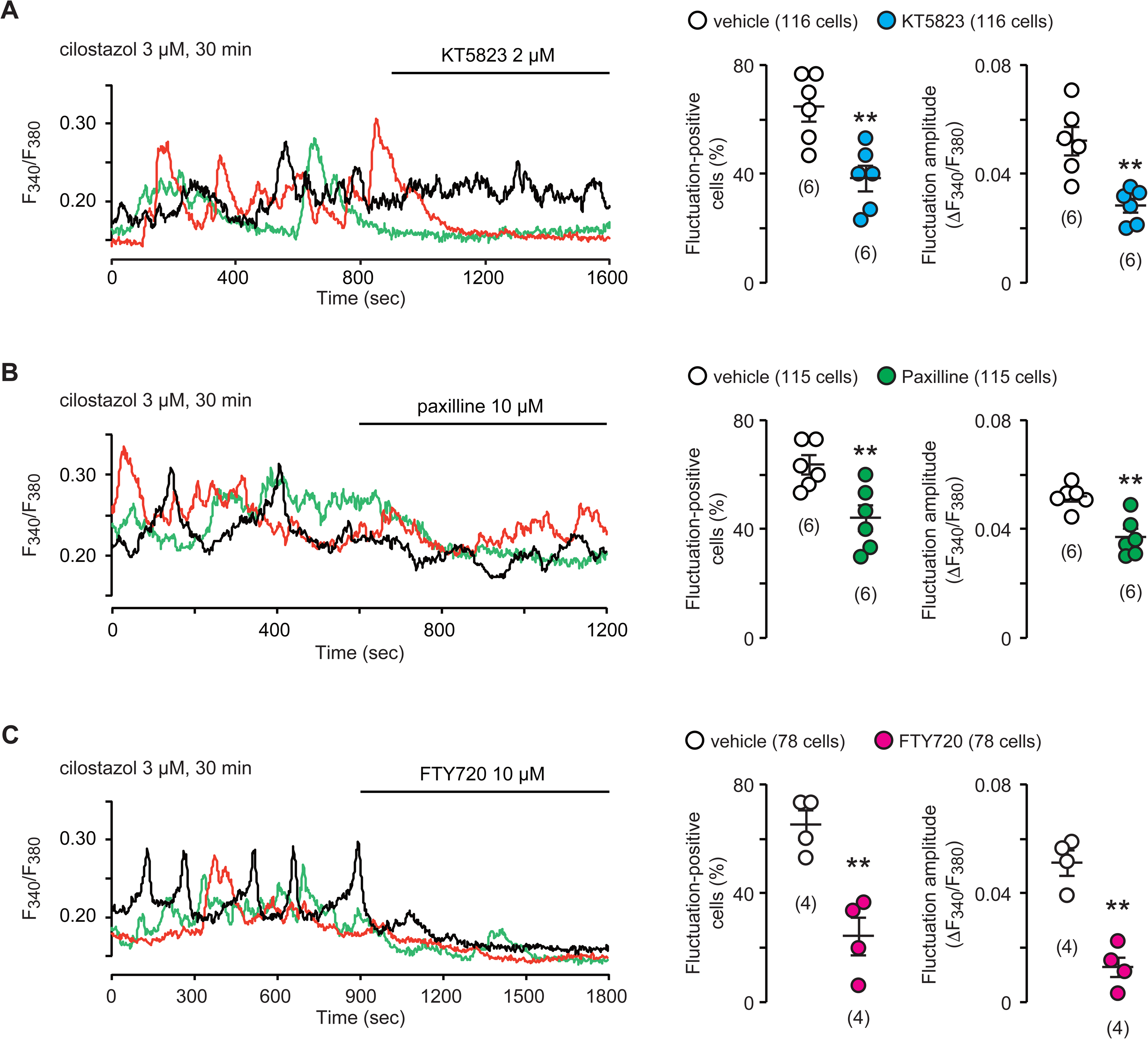
Pharmacological characterization of cilostazol-facilitated Ca^2+^ fluctuations. **(A)** Effects of the PKG inhibitor KT5823 on cilostazol-facilitated Ca^2+^ fluctuations. Femoral bone slices were pretreated with cilostazol for 30 min and then subjected to Ca^2+^ imaging. Representative recording traces from three round chondrocytes are shown (left panel), and KT5823-evoked effects on Ca^2+^ fluctuations were summarized (right graphs). **(B)** Effects of the BK channel inhibitor paxilline on cilostazol-facilitated Ca^2+^ fluctuations. Representative recording traces (left panel) and a summary of paxilline-evoked effects are shown (right graphs). **(C)** Effects of the TRPM7 inhibitor FTY720 on cilostazol-facilitated Ca^2+^ fluctuations. Representative recording traces (left panel) and a summary of FTY720-evoked effects (right graphs) are shown. The data are presented as the mean ± SEM., and the numbers of mice and cells examined are shown in the bar graph and keys, respectively. Significant inhibitor-induced shifts are marked with asterisks (\*\**p* < 0.01 in t-test).

### Cilostazol reduces the membrane potential of growth plate chondrocytes

Next, growth plate slices were subjected to confocal imaging using the voltage-dependent dye oxonol VI (Yamazaki *et al*, 2011). In this measurement, depolarization results in the accumulation of the dye in cells and thus linearly correlates well with the cellular fractional fluorescence intensity which is normalized to the maximum intensity monitored using a 100 mM KCl-containing bath solution. Under resting conditions, cilostazol pretreatment (3 μM for 30 min) reduced the fractional intensities in growth plate chondrocytes (Fig. 6A); the estimated resting potentials were -40.0 ± 0.2 and -41.3 ± 0.1 mV in the vehicle- and cilostazol-pretreated round chondrocytes, respectively.

**Fig. 6.**
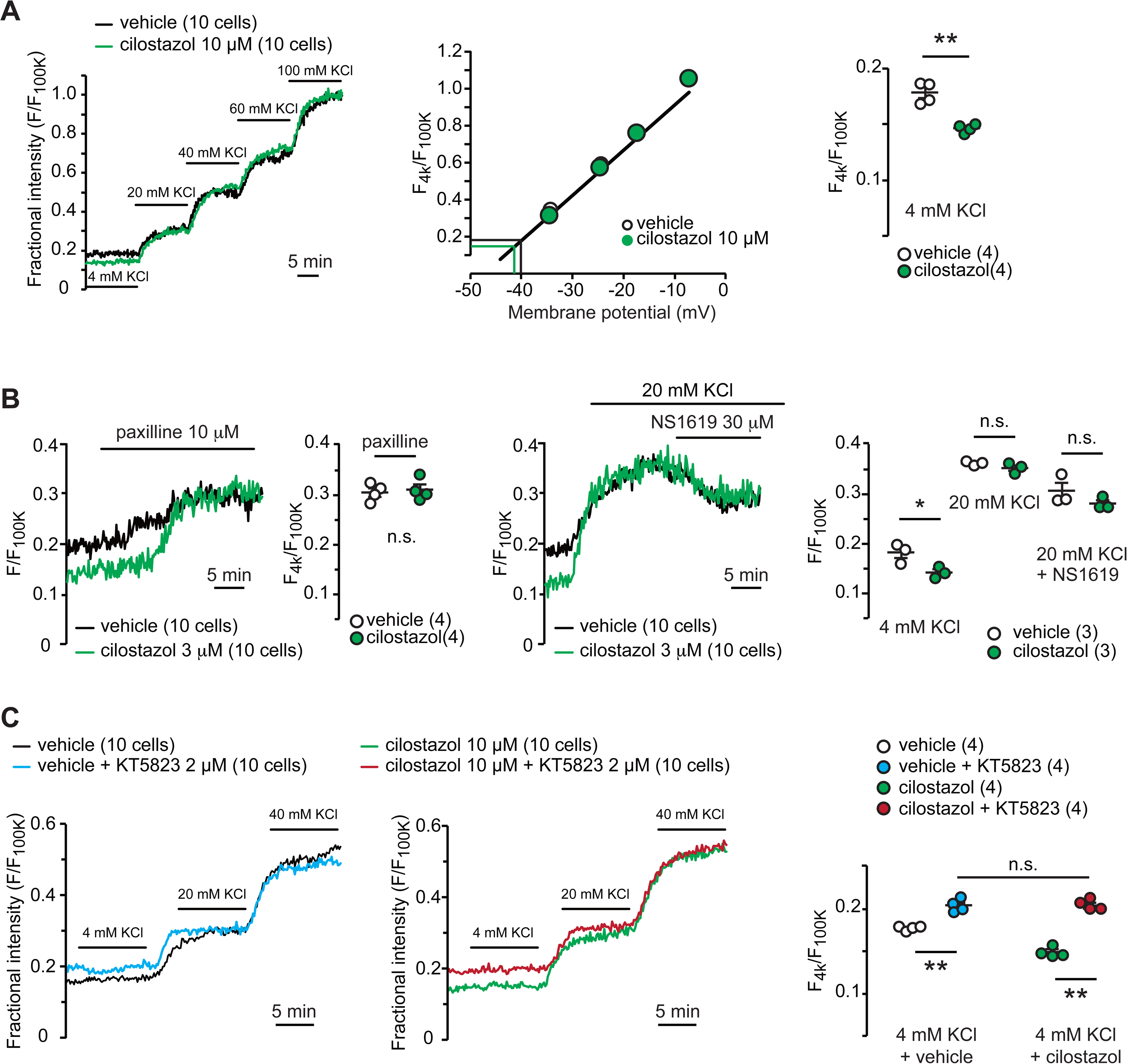
Cilostazol decreases the membrane potential of growth plate chondrocytes. **(A)** Confocal oxonol VI imaging of round chondrocytes. Femoral bone slices were pretreated with or without cilostazol for 30 min and then subjected to membrane potential imaging. During contiguous perfusion of high-K^+^ solutions, cellular fluorescence intensities were monitored and normalized to the maximum value in the 100 mM KCl-containing solution to yield the fractional intensity. Representative averaged traces from ten cells are shown (left panel). For preparation of the calibration plot (middle graph), the data from 10 cells in bathing solutions containing 4 (normal solution), 20, 40, 60, and 100 mM KCl are summarized; green and black lines indicate the estimated resting membrane potentials of the cilostazol- and vehicle-pretreated cells, respectively. The resting fractional intensities were quantified and statistically analyzed in the cilostazol- and vehicle-pretreated cells (left graph). The data are presented as the mean ± SEM., and the numbers of mice examined are shown in parentheses in the keys. Significant differences between the cilostazol- and vehicle-pretreated cells are indicated by asterisks (\*\**p* < 0.01 according to t test). **(B)** Effects of BK channel modulators on the resting membrane potential in round chondrocytes. The BK channel inhibitor paxilline abolished the difference between the vehicle- and cilostazol-treated cells (left panel and graph). The recording data from ten cells pretreated with or without cilostazol were averaged and the fractional intensities elevated by paxilline are summarized. The BK channel activator NS1619 canceled the difference between vehicle- and cilostazol-treated cells (right panel and graph). Recording data from ten cells pretreated with or without cilostazol were averaged, and the fractional intensities in the normal, 20 mM KCl and NS1619-containing 20 mM KCl solutions are summarized. The data are presented as the mean ± SEM., the numbers of mice examined are shown in parentheses in the keys and significant differences between the cilostazol- and vehicle-pretreated cells are indicated by asterisks (\**p* < 0.05 according to t test). **(C)** Effect of the PKG inhibitor KT5823 on the resting membrane potential in round chondrocytes. After pretreatment with or without KT5823 and cilostazol, round chondrocytes in femoral bone slices were examined. In each pretreated group, the recording data of ten cells were averaged (left panels), and the resting fractional intensities are summarized (right graph). The data are presented as the mean ± SEM., and the numbers of mice examined are shown in parentheses. Significant differences between each group are indicated by asterisks (\*\**p* < 0.01 according to two-way ANOVA and Sidak’s test).

However, the addition of paxilline to a perfusion solution quickly elevated the fractional intensities and abolished the difference between the cilostazol- and vehicle-pretreated cells (Fig. 6B). In the presence of the BK channel activator NS1619, the cilostazol- and vehicle-pretreated cells exhibited similar fractional intensities. Therefore, cilostazol likely activated BK channels to induce hyperpolarization.

In the CNP-evoked signaling cascade, PKG phosphorylates and activates BK channels (Fig. EV1). Even in the vehicle-pretreated control chondrocytes, pretreatment with KT5823 clearly elevated the resting membrane potential, suggesting that the mechanism of PKG-mediated BK channel activation was functioning under basal conditions (Fig. 6C). Furthermore, the shift induced by KT5823 cotreatment was more obvious in the cilostazol-pretreated cells, and KT5823 seemed to abolish the difference in membrane potential between the vehicle- and cilostazol-pretreated cells. Therefore, cilostazol likely enforced the PKG-mediated BK channel activation in growth plate chondrocytes.

### Cilostazol activates CaMKII in growth plate chondrocytes

CNP activates TRPM7-mediated Ca^2+^ entry and thus stimulates CaMKII to facilitate ECM formation in growth plate chondrocytes (Fleig & Chubanov, 2014; Miyazaki *et al*., 2022). Immunohistochemical analysis of round chondrocytes using an antibody against autophosphorylated CaMKII (pThr 286) revealed that the cilostazol-treated cells (3 μM for 30 min) were decorated more frequently and densely than the control cells (Fig. 7A). The CaMKII inhibitor KN93 totally abolished the immunodecoration enhanced by cilostazol, indicating that the antibody precisely recognized the phosphorylated antigen. Furthermore, Western blot analysis of cell lysates prepared from femoral growth plates detected that the cilostazol treatment significantly increased the phospho-CaMKII content without affecting the total CaMKII density (Fig. 7B). Therefore, cilostazol seemed to activate CaMKII downstream of facilitated TRPM7-mediated Ca^2+^ entry in growth plate chondrocytes. Taken together with all the pharmacological data above, PDE3 inhibitors seemed to stimulate the cGMP-driven PKG-BK channel-TRPM7 channel-CaMKII signaling axis in the same manner as CNP.

**Fig. 7.**
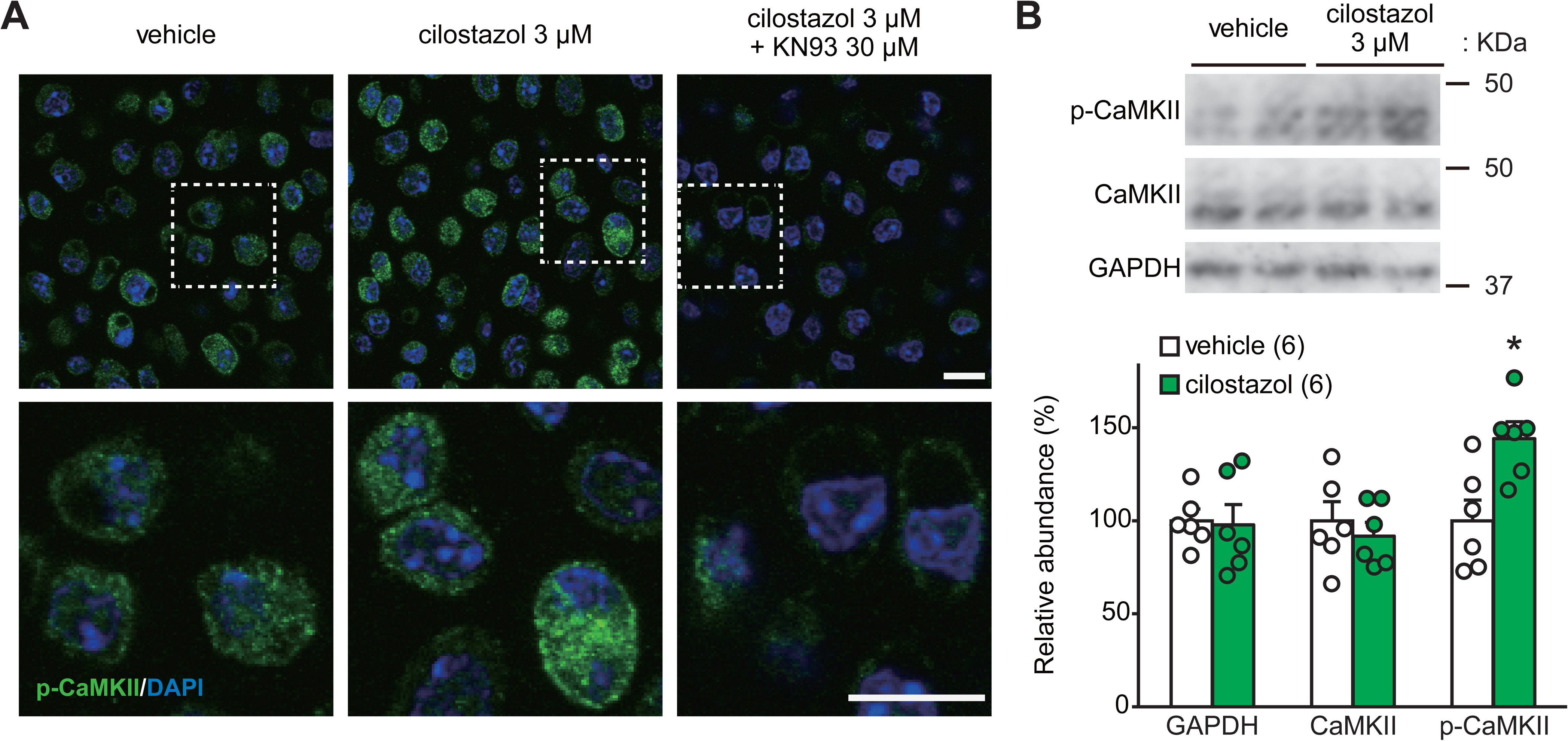
CaMKII activation in cilostazol-treated chondrocytes. **(A)** Immunohistochemical staining for phospho-CaMKII (p-CaMKII) in round chondrocytes. Femoral bone slices were pretreated with or without cilostazol and the CaMKII inhibitor KN93, and then subjected to immunostaining with an antibody against p-CaMKII. 4′,6-Diamidino-2-phenylindole (DAPI) was used for nuclear staining. The lower panels show high-magnification images of the white-dotted regions in the upper panels. Scale bars, 10 μm. **(B)** Immunoblot analysis of CaMKII in growth plate cartilage. Growth plate lysates were prepared from femoral bone slices pretreated with or without cilostazol and subjected to immunoblot analysis with antibodies against total CaMKII and p-CaMKII (upper panel). Glyceraldehyde-3-phosphate dehydrogenase (GAPDH) was also analyzed as a loading control. The immunoreactivities observed are summarized in the lower graph. The data are presented as the mean ± SEM., and the numbers of mice examined are shown in parentheses. A significant difference between the cilostazol- and vehicle-pretreated cells is indicated by an asterisk (\**p* < 0.05 according to t test).

### Combined effects of CNP and cilostazol on cultured bone growth

We finally examined the combined effects of CNP and cilostazol on cultured bone elongation. Moderate (10 nM) and submaximal (30 nM) CNP concentrations resulted in increased of 127.6 ± 0.6% and 129.8 ± 4.1%, respectively, in the longitudinal size of metatarsal bones during 4 days of culture. Cilostazol (10 μM) further stimulated bone outgrowth in 10 nM CNP-containing medium (Fig. 8A) but had no effect on bone outgrowth in 30 nM CNP-containing medium (Fig. 8B). Therefore, cilostazol seemed to lose its pharmacological effect under high-CNP conditions. These observations were consistent with our conclusion that CNP and cilostazol stimulated the common signaling axis to promote bone growth.

**Fig. 8.**
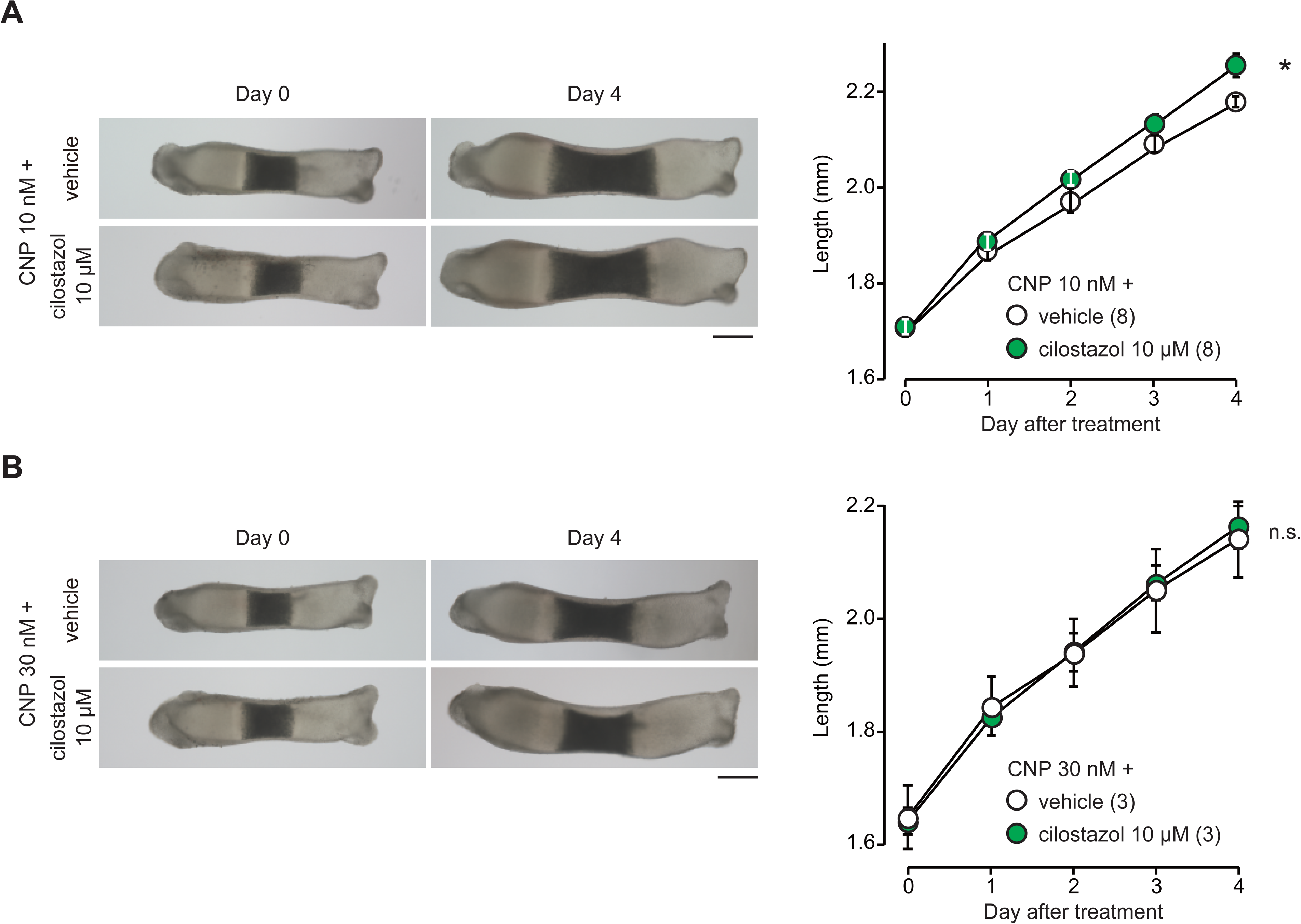
Combinational effects of CNP and cilostazol on cultured bone outgrowth. (**A**) Combinational effects of 10 nM CNP and 10 μM cilostazol. (**B**) Combinational effects of 30 nM CNP and 10 μM cilostazol. Representative images of the metatarsals before and after culture for 4 days under the indicated conditions are shown (left panels; scale bar, 0.3 mm). Longitudinal growth of the metatarsals was quantified over the culture period (right graphs). The numbers of mice examined are shown in parentheses. The bone sizes at the endpoint were analyzed and a statistical difference from the vehicle-treated group is marked with an asterisk (\**p* < 0.05; n.s., not significant according to two-way ANOVA and Sidak’s test).

## DISCUSSION

In this study, we demonstrated that PDE3 inhibitors facilitate cultured bone outgrowth (Fig. 2) and increase body size in juvenile mice (Fig. 3). Our observations also suggest that cilostazol elevates cGMP levels, activates BK channels via PKG-catalyzed phosphorylation, and stimulates TRPM7-mediated Ca^2+^ entry by increasing the ion driving force (Fig. EV 4-6), thus facilitating CaMKII (Fig.7) in growth plate chondrocytes. Therefore, PDE3 inhibitors probably enforce the CNP signaling cascade (Fig. EV1) and seem to stimulate ECM secretion downstream of CaMKII activation, leading to facilitating bone elongation. This conclusion is further supported by the data that cilostazol did not stimulate the outgrowth of mutant bones from chondrocyte-specific *Trpm7*-knockout mice (Fig. EV4B). In standard clinical treatments, cilostazol is used as an oral anti-platelet aggregatory drug, while olprinone and milrinone are intravenously administered as inotropic and vasodilative drugs (Kherallah *et al*, 2021; Young & Ward, 1988). Our data may predict that PDE3 inhibitors would be applicable for the treatment of various types of syndromes involving short statue via drug repositioning. The CNP variant vosoritide has been clinically used via daily subcutaneous injection for achondroplasia treatment (Legeai-Mallet, 2016; Savarirayan *et al*., 2020). Indication expansion trials using approved PDE3 inhibitors might provide patient-friendly oral drugs with economic benefits to combat various syndromes characterized by short stature. Moreover, based on the data from combined CNP and cilostazol treatments in bone culture (Fig. 8), PDE3 inhibitors can be used in new combination therapies with vosoritide for achondroplasia patients.

The gene expression profile likely suggests that several PDE subtypes differentially contribute to the inactivation of cGMP and cAMP in growth plate chondrocytes (Fig. 1). PDE3B seems to mainly maintain the intracellular cGMP concentration under resting conditions, because cilostazol and milrinone elevated the steady-state cGMP concentration in growth plate specimens (Fig. EV 4 and EV6). From a biochemical perspective, it is interesting to address the subtype-specific roles of the PDE species expressed in growth plate chondrocytes. Among the PDE species presumed to be expressed in the chondrocytes, PDE1B, PDE2A, PDE3B, PDE6G and PDE10A might differentially become active to hydrolyze cGMP and cAMP in response to various cellular conditions (Heckman *et al*., 2018),, and thus could modulate both the cGMP and cAMP signaling cascades. It has been reported that cAMP-dependent protein kinase (PKA), together with PKG, phosphorylates and activates BK channels in several cell types (Calderone, 2002; Kyle & Braun, 2014). Therefore, it is reasonably hypothesized that the CNP signaling cascade might be cooperatively controlled by PKG and PKA (Fig. EV1), both of which may be complexly regulated by the PDE subtypes in growth plate chondrocytes. To better understand the signal crosstalk proposed in CNP-induced long-bone outgrowth, the roles of the PKA and PDE subtypes would be further focused in developing growth plates.

## MATERIALS AND METHODS

### Reagents, primers and mice

The commercial reagents used in this study are listed in Table EV1. The synthetic primers used for RT-PCR analysis and mouse genotyping are listed in Table EV2. C57BL/6 mice were supplied by Japan SLC Inc. Chondrocyte-specific *Trpm7* knockout mice were generated and genotyped as described previously (Miyazaki *et al*., 2022). Achondroplasia model mice (*Fgfr3*^ach^ mice) were generated as described in a previous report (Naski *et al*, 1998): the activated fibroblast growth factor receptor 3 (FGFR3) transgene is driven by the *Col2a1* promoter and enhancer sequences in *Fgfr3*^ach^ mice (FVB/N background). All experiments in this study were conducted with the approval of the Animal Research Committee according to the regulations on animal experimentation at Kyoto University (approval no. 17-7-5).

### Gene expression analysis

For gene chip analysis, distal epiphyses enriched in round chondrocytes were separated from embryonic day 17.5 (E17.5) femoral bones under a stereomicroscope and subjected to total RNA preparation using the commercial reagent Isogen (Nippon Gene Co., Japan). The resulting total RNA was reverse-transcribed and analyzed using the GeneChip Mouse Genome 430 2.0 (Affymetrix) according to the manufacturer’s instructions. The obtained data have been deposited in the National Center for Biotechnology Information-Gene Expression Omnibus (NCBI-GEO) database under accession number GSE105256. For quantitative RT-PCR analysis, the total RNA was reverse-transcribed using a commercial kit (ReverTra ACE qPCR-RT kit). The resulting cDNAs were analyzed by real-time PCR (LightCycler 480 II, Roche), and the cycle threshold was determined from the amplification curve as an index for relative mRNA content in each reaction.

### Metatarsal organ culture

Metatarsal bone rudiments were cultured as previously described (Staines *et al*., 2019). Briefly, the central metatarsal rudiments were dissected from E15.5 mice and cultured in αMEM supplemented with 5 μg/mL ascorbic acid, 1 mM β-glycerophosphate pentahydrate, 100 units/mL penicillin, 100 μg/mL streptomycin and 0.2% bovine serum albumin (fatty acid free). The explants were analyzed under a photomicroscope (BZ-X710, Keyence, Japan) for size measurements using Fiji/ImageJ software (US. NIH). For histological analysis, the cultured bones were fixed in 4% paraformaldehyde, embedded in Super cryoembedding medium (Section-lab, Japan), and frozen in liquid nitrogen. Serial cryosections (∼6 μm in thickness) were prepared and treated with a commercial hematoxylin and eosin solution (Wako Pure Chemical, Japan) or alcian blue solution (Merck, USA) for photomicroscopic observation and image analysis.

### In vivo assessment of cilostazol-injected effects

C57BL/6 mice (3 weeks old) were intraperitoneally (i.p.) injected daily with 10 mg/kg/day of cilostazol suspended in olive oil (Wako, Japan) for 4 weeks. The vehicle-treated mice were injected with olive oil. The naso-anal length and body weight were measured weekly as previously reported (Ueda *et al*, 2019); the cranium was fixed and the mouse body was fully stretched under anesthesia for the measurement.

After the test period, the heart rate and blood pressure of the mice were measured using a tail-cuff sphygmomanometer (BP-98AL, Softron, Japan). Individual blood was then drawn from the inferior vena cava of mouse under anesthesia and centrifuged at 1500 × g for 15 min to collect the serum. Serum biochemical data for Na^+^, K^+^, Cl^-^, Ca^2+^, aspartate aminotransferase (AST), lactate dehydrogenase (LDH) and creatinine kinase (CK) were obtained by Oriental Yeast Co. (Nagahama, Japan). Finally, the femoral and tibial bones were isolated from the mice and fixed in 4% paraformaldehyde, demineralized in 18 w/v% EDTA solution (pH 7.2) for two weeks, embedded in Super cryoembedding medium (Section-lab, Japan), and frozen in liquid nitrogen. Serial cryosections (∼8 μm in thickness) were prepared and treated with a commercial alcian blue solution (Merck, USA). Microscopic images were quantitatively analyzed using Fiji/ImageJ software. The average cell density and alcian blue positive area were calculated from three independent visual fields (300 x 600 μm) in each growth plate preparation.

### Bone slice preparations

Femoral bones isolated from E17.5 mice were immersed in a physiological salt solution (PSS): (in mM) 150 NaCl, 4 KCl, 1 MgCl_2_, 2 CaCl_2_, 5.6 glucose, and 5 HEPES (pH 7.4). Longitudinal bone slices (40-100 µm thickness) were prepared using a vibrating microslicer (DTK-1000N, Dosaka EM Co., Japan) as previously described (Miyazaki *et al*., 2022) and subjected to cyclic nucleotide measurements, Ca^2+^ imaging, membrane potential monitoring, and immunochemical analysis.

### Quantification of cAMP and cGMP

Epiphyseal specimens enriched in proliferating chondrocytes were removed from the femoral bone slices under stereomicroscope and treated with or without PDE3 inhibitors (3 μM cilostazol or 10 μM milrinone for 30 min) in PSS at 37 °C. For cGMP quantification, the specimens (8-10 slices) were homogenized in 5% trichloroacetic acid (TCA, 0.3 mL), and for cAMP quantification, the specimens (4-5 slices) were homogenized in 5% TCA (0.6 mL) by a microhomogenizer (NS-310E, MICROTEC, Japan). The lysates were then centrifuged at 1500 × g for 10 min to remove insoluble materials. The precipitates were recovered for protein quantification (#23225, Pierce BCA assay, Thermo Fisher Scientific), and the supernatants were subjected to cAMP and cGMP quantification using commercial kits (ADI-900-066A cAMP ELISA kit and ADI-900–014 cGMP ELISA kit, Enzo Life Sciences) after removing TCA by ether extraction three times.

### Ca^2+^ imaging

Fura-2 Ca^2+^ imaging of bone slices was performed as previously described (Ichimura *et al*, 2023; Miyazaki *et al*., 2022). Briefly, bone slices placed on glass-bottom dishes (Matsunami, Japan) were incubated in PSS supplemented with 15 μM Fura-2AM for 1 hr at 37 °C. Fluorescence microscopy distinguished round, columnar, and hypertrophic chondrocytes with characteristic morphological features in the bone slices loaded with Fura-2. For ratiometric imaging, excitation light at 340 and 380 nm was alternately delivered, and emission light at >510 nm was detected by a cooled EM-CCD camera (Model C9100-13; Hamamatsu Photonics, Japan) mounted on an upright fluorescence microscope (DM6 FS, Leica) using a 40x water-immersion objective (HCX APO L, Leica). In typical measurements, ∼30 round chondrocytes were randomly examined in each slice preparation to select the Ca^2+^-fluctuation-positive cells generating spontaneous events (>0.025 in Fura-2 ratio) using commercial software (Leica Application Suite X), and recording traces from the positive cells were then analyzed using Fiji/ImageJ software for examining Ca^2+^ fluctuation amplitude and frequency. Imaging experiments were performed at room temperature (∼25 °C), and PSS was used as the normal bathing solution. For the pretreatments of each compound, bone slices were immersed in PSS with the indicated compound for 30 min at 37 °C after Fura-2 loading.

### Membrane potential monitoring

Membrane potential recording was performed as previously described (Miyazaki *et al*., 2022). Briefly, bone slices placed on glass-bottom dishes were perfused with PSS containing 200 nM oxonol VI (AnaSpec, USA) for 45 min at room temperature (∼25 °C). During the recording, hyperpolarization results in extrusion of the dye from cells to subsequently decrease the cellular fluorescence intensity, while depolarization increases the fluorescence intensity. For preparation of the calibration plot showing the relationship between the fluorescence intensity and membrane potential, PSS solutions containing 20 mM, 40 mM, 60 mM or 100 mM KCl were used as bathing solutions. Fluorescence images with excitation at 559 nm and emission at >606 nm were captured at a sampling rate of ∼10 s using a confocal microscope system (FV1000, Olympus, Japan). For the pretreatments with each compound, bone slices were immersed in PSS with the indicated compound for 30 min at 37 °C before oxonol VI loading.

### Immunochemical analysis

For immunohistochemical analysis, bone slices were pretreated with or without cilostazol and fixed in 4% paraformaldehyde. After treatment with 1% hyaluronidase to increase immunodetection (Ahrens & Dudley, 2011; Mouser *et al*, 2016) and blocking with fetal bovine serum-containing solution, the bone slices were incubated with primary and Alexa 488-conjugated secondary antibodies and observed with a confocal microscope (FV1000, Olympus, Japan). For immunoblot analysis, bone slices were lysed in the buffer containing 4% sodium deoxycholate, 20 mM Tris-HCl (pH 8.8) and a phosphatase inhibitor cocktail (100 mM NaF, 10 mM Na_3_PO_4_, 1 mM Na_2_VO_3_ and 20 mM β-glycerophosphate). The resulting lysate proteins were electrophoresed on SDS-polyacrylamide gels and electroblotted onto nylon membranes for immunodetection using primary and HRP-conjugated secondary antibodies. The antigen proteins were visualized using a chemiluminescence reagent and image analyzer (Amersham Imager 600, Cytiva). The immunoreactivities yielded were quantitatively analyzed using Fiji/ImageJ software.

### Quantification and statistical analysis

All the data obtained are presented as the means ± SEMs. with *n* values indicating the number of examined mice. Student’s t test and ANOVA were used for two-group and multiple group comparisons, respectively (Prism 7, GraphPad Software Inc.): *p* < 0.05 was considered to indicate statistical significance.

## Data availability

The datasets produced in this study are available in the following databases: Microarray data data: Gene Expression Omnibus GSE105256 (https://www.ncbi.nlm.nih.gov/geo/query/acc.cgi?acc=GSE105256)

## Author contribution

T.K. and A.I. are equally contributing first authors. T.K. and A.I. conducted Ca^2+^ imaging. T.K., A.I., T.Y., J.L., G.E.K. and H.I. conducted biochemical and cell-physiological analyses. Y.U. maintained and provided the *Fgfr3*-Tg mice. A.I. and H.T. oversaw the project and drafted the manuscript. All authors reviewed/edited the manuscript.

## Acknowledgments

We thank Jun Matsushita (Graduate School of Pharmaceutical Sciences, Kyoto University) for mouse reproduction by *in vitro* fertilization. This work was supported in part by the MEXT/JSPS (KAKENHI 21H02663, 21K19565, 23H02687 and 23K08687), Platform Project for Supporting Drug Discovery and Life Science Research (22ama121034j0001), Takeda Science Foundation, Kobayashi International Scholarship Foundation, the NAKATOMI Foundation, the Mother and Child Health Foundation and Japan Foundation for Applied Enzymology. The research activity of T.K. is supported by the JST SPRING program (JPMJSP2110).

---

**Fig. EV1. CNP signaling pathway and PDE3 inhibitors.**

CNP stimulates the cGMP-driven PKG-BK channel–TRPM7 channel–CaMKII axis in growth plate chondrocytes and thus promotes bone growth by facilitating ECM synthesis (Miyazaki et al., 2022). Our present study suggested that PDE3 inhibitors facilitate the signaling axis by inhibiting cGMP hydrolysis.

**Fig. EV2. Effects of various PDE inhibitors on cultured bone growth.**

The effects of anagrelide (**A**), olprinone (**B**), BAY-60-7550 (**C**) and MP 10 (**D**) on cultured metatarsal growth were examined. Representative images of the metatarsals before and after 4 days of culture in the presence or absence of the PDE inhibitors (left panels). Scale bar, 0.3 mm. Longitudinal bone growth cultured with or without the PDE inhibitors was quantified over the culture period (right graphs). The numbers of mice examined are shown in parentheses. The bone sizes were analyzed in each experiment, and statistical differences at the endpoint from the vehicle-pretreated group are marked with asterisks (\*\**p* < 0.01 according to two-way analysis of variance (ANOVA) and Sidak’s test).

**Fig. EV3. Milrinone promoted cultured bone growth.**

**(A)** Milrinone-promoted outgrowth of cultured metatarsal bones. Representative images of the metatarsals before and after culture for 4 days in the presence or absence of milrinone (left panels; scale bar, 0.3 mm). Longitudinal growth of the metatarsals was quantified over the culture period (right graph). The numbers of mice examined are shown in parentheses. The bone lengths were analyzed at the endpoint, and statistical difference from the vehicle-treated group is marked with an asterisks (\**p*<0.05 according to two-way ANOVA and Sidak’s test). **(B)** Histological analysis of metatarsal bones cultured in the presence or absence of milrinone. The mid-longitudinal sections of cultured metatarsal bones were stained with hematoxylin and eosin (upper left panels; scale bar, 0.3 mm), and their high-magnification images from the round (R), columnar (C), and hypertrophic (H) chondrocyte zones are also shown (scale bar, 30 μm). The metatarsal sections were likewise stained with alcian blue (upper right panels; scale bar, 0.3 mm). MD indicates the mid-diaphysis. In the round, columnar and hypertrophic chondrocyte zones, the zonal size, cell density and alcian blue-positive area were examined and are summarized (lower graphs). The data are presented as the mean ± SEM., and the numbers of mice examined are shown in parentheses. Statistical differences between the vehicle- and cilostazol-treated groups are marked with asterisks (\**p* < 0.05 and \*\**p* < 0.01 according to t test).

**Fig. EV4. Effects of cilostazol on the mutant bone growth.**

**(A)** Effects of cilostazol on the outgrowth of cultured metatarsals from achondroplasia model mice. Metatarsals isolated from control (wild-type) and achondroplasia model embryos (*Fgfr3*-Tg) were precultured in regular medium for 5 days, and then test-cultured in medium supplemented with or without cilostazol for 4 days. Representative images of cultured metatarsals are shown (left panels; scale bar, 0.3 mm). Longitudinal bone outgrowth during the test period was statistically analyzed in each genotype group (right graph). The numbers of mice examined are shown in parentheses. The bone sizes were analyzed over the test period and statistical differences between the cilostazol-treated and nontreated groups for each genotype are marked with asterisks (\**p* < 0.05 according to two-way ANOVA and Sidak’s test). **(B)** Cilostazol had no effect on the outgrowth of *Trpm7*-deficient bones. Metatarsals isolated from chondrocyte-specific *Trpm7*-knockout (*Trpm7*^fl/fl^, *Col11Enh-Cre*^+/−^) embryos were precultured in regular medium for 5 days, and then test-cultured in medium supplemented with or without cilostazol for 4 days. Representative images of the metatarsals before and after the test are shown (left panels; scale bar, 0.3 mm), and the longitudinal growth of the metatarsals was examined over the test culture period (right graph). The numbers of mice examined are shown in parentheses, and the data obtained were statistically analyzed (two-way ANOVA and Sidak’s test; n.s., not significant).

**Fig. EV5. No obvious effects of *in vivo* cilostazol treatments on major cardiovascular and serological parameters.**

After daily injection of cilostazol for 4 weeks, 7-week-old mice were examined. **(A)** Heart rate and blood pressure were measured by the tail-cuff method. **(B)** Mouse serum was prepared from blood drawn from the inferior vena cava and subjected to outsourced serological analyses. The data are presented as the mean ± SEM., and the numbers of mice examined are shown in parentheses. The data were statistically analyzed, and no significant difference was observed between the vehicle- and cilostazol-treated groups (t test).

**Fig. EV6. Milrinone. Cilostazol facilitates Ca^2+^ fluctuations in growth plate chondrocytes.**

**(A)** Effects of milrinone on cGMP and cAMP levels in growth plate chondrocytes. Femoral bone slices enriched in growth plate chondrocytes were treated with or without milrinone for 30 min and then subjected to cyclic nucleotide quantification. The data are presented as the mean ± SEM., and the numbers of mice examined are shown in parentheses. A significant difference between the groups is indicated by asterisks (“\*\**p* < 0.01 according to t test). **(B)** Effects of milrinone on Ca^2+^ fluctuations in in growth plate chondrocytes. Fura-2 imaging of round chondrocytes in femoral bone slices pretreated with or without cilostazol for 30 min. Representative recording traces from three cells are shown for each pretreatment group (upper panels). The effects of milrinone on spontaneous Ca^2+^ fluctuations are summarized (lower graphs); the fluctuation-positive cell ratio, fluctuation amplitude and fluctuation frequency were statistically analyzed. The numbers of mice and cells examined are shown in the bar graphs and keys, respectively. Statistical differences from the vehicle-pretreated group are marked with asterisks (\**p* < 0.05 and ** *p* < 0.01 according to t test).

**Table EV1:** Reagents used in this study.

| Reagents | supplier | identifier | dilution |
| --- | --- | --- | --- |
| Anti-phospho-CaMKII (Thr286)<br>(Rabbit monoclonal) | Cell Signaling Technology | Cat#12716; RRID: AB_271389 | WB (1/1000),<br>IF (1/200) |
| Anti-CaMKII<br>(Rabbit monoclonal) | Abcam | Cat#EP1829Y; RRID: AB_868841 | WB (1/1000) |
| Anti-GAPDH<br>(Rabbit polyclonal) | Sigma-Aldrich | Cat#G9545; RRID: AB_796208 | WB (1/10000) |
| Anti-rabbit IgG-HRP<br>(Mouse monoclonal) | Santa Cruz | Cat#sc-2357; RRID: AB_628497 | WB (1/2000) |
| Anti-rabbit Alexa Fluor 488<br>(Goat polyclonal) | Invitrogen | Cat#A-11008; RRID: AB_143165 | IF (1/500) |
| NaCl | Wako | Cat#191-01665 |  |
| KCl | Wako | Cat#163-03545 |  |
| CaCl<sub>2</sub> · 2H<sub>2</sub>O | Wako | Cat#033-25035 |  |
| MgCl<sub>2</sub> · 6H<sub>2</sub>O | Wako | Cat#135-00165 |  |
| Glucose | Wako | Cat#049-31165 |  |
| Glycine | Wako | Cat#077-00735 |  |
| 2-Amino-2-hydroxymethyl-1,3-propanediol (Tris) | Nacalai tesque | Cat#35408-91 |  |
| 2-Amino-2-Hydroxymethyl-1,3-propanediol hydrochloride (Tris-HCl) | Wako | Cat#012-17455 |  |
| Olive oil | Wako | Cat#150-00276 |  |
| HEPES | Dojindo | Cat#342-01375 |  |
| Methanol | Wako | Cat#139-01827 |  |
| Ethanol | Junsei | Cat#17065-0382 |  |
| Diethyl ether | Wako | Cat#052-01165 |  |
| Chloroform | Nacalai tesque | Cat#08402-55 |  |
| 2-propanol | Nacalai tesque | Cat#29113-95 |  |
| EDTA · 2Na | Dojindo | Cat#345-01865 |  |
| Trichloroacetic acid | Nacalai tesque | Cat#34603-02 |  |
| L-ascorbic acid 2-phosphate<br>sesquimagnesium salt hydrate | Nacalai tesque | Cat#13570-82 |  |
| β-glycerophosphate $\cdot 5H_2O$ | Wako | Cat#048-34332 | |
| Penicillin-streptomycin solution | Wako | Cat#168-23191 |  |
| paraformaldehyde | Nacalai tesque | Cat#09154-85 |  |
| Mayer's hematoxylin solution | Wako | Cat#131-09665 |  |
| 1 % eosin Y solution | Wako | Cat#051-06515 |  |
| Alcian-blue solution pH2.5 | Merck Millipore | Cat#101647 |  |
| Super cryoembedding medium<br>(SCEM) | Section-Lab |  |  |
| MEM $\alpha$ | Thermo Fisher Scientific | Cat#12561056 | |
| Albumin solution from bovine<br>serum (fatty acid free) | Wako | Cat#017-22231 |  |
| Cilostazol | Tokyo Chemical Industry | Cat#C2587 |  |
| Milrinone | Tokyo Chemical Industry | Cat#M1663 |  |
| Anagrelide hydrochloride | Cayman | Cat#21411 |  |
| Olprinone hydrochloride | MedChemExpress | Cat#H-14254 |  |
| BAY-60-7550 | Cayman | Cat#10011135 |  |
| MP10 | Cayman | Cat#18266 |  |
| Fura-2 AM special packaging | Dojindo | Cat#F025 |  |
| Oxonol VI | AnaSpec | Cat#AS-84704 |  |
| FTY720 | Sigma-Aldrich | Cat#SML0700 |  |
| Paxilline | Tocris Bioscience | Cat#2006/10 |  |
| KT5823 | Cayman | Cat#10010965 |  |
| KN-93 | Wako | Cat#115-00641 |  |
| Fetal bovine serum | Wako | Cat#533-69545 |  |
| Hyaluronidase from sheep testes | Sigma-Aldrich | Cat#H2126 |  |
| Sodium deoxycholate | Wako | Cat#192-08312 |  |
| NaF | Nacalai tesque | Cat#31420-82 |  |
| $Na_3PO_4 \cdot 12H_2O$ | Nacalai tesque | Cat#31804-75 | |
| $Na_3VO_4$ | Wako | Cat#198-09752 | |
| Protease Inhibitor Cocktail for Use with Mammalian Cell and Tissue Extracts | Nacalai tesque | Cat#25955-24 |  |
| Sodium dodecyl sulfate | Wako | Cat#194-13985 |  |
| Acrylamide/Bis mixed solution | Nacalai tesque | Cat#06141-35 |  |
| WIDE RANGE Gel Preparation Buffer(4x) for PAGE | Nacalai tesque | Cat#07831-94 |  |
| TEMED | Nacalai tesque | Cat#33401-72 |  |
| Ammonium Peroxodisulfate | Wako | Cat#012-20503 |  |
| Blocking One | Nacalai tesque | Cat#03953-95 |  |
| ISOGEN | Nippon Gene | Cat#319-90211 |  |
| ReverTra Ace qPCR RT Master Mix with gDNA Remover | Toyobo | Cat#FSQ-301 |  |
| FastStart Essential DNA Green Master Mix | Roche | Cat#06924204001 |  |
| KOD FX | Toyobo | Cat#KFX-101X5 |  |
| Pierce BCA protein assay kit | Thermo Fisher Scientific | Cat#0023225 |  |
| cAMP ELISA kit | Enzo Life Sciences | Cat#ADI-900-066A |  |
| cGMP ELISA kit | Enzo Life Sciences | Cat#ADI-900-014 |  |

**Table EV2:**
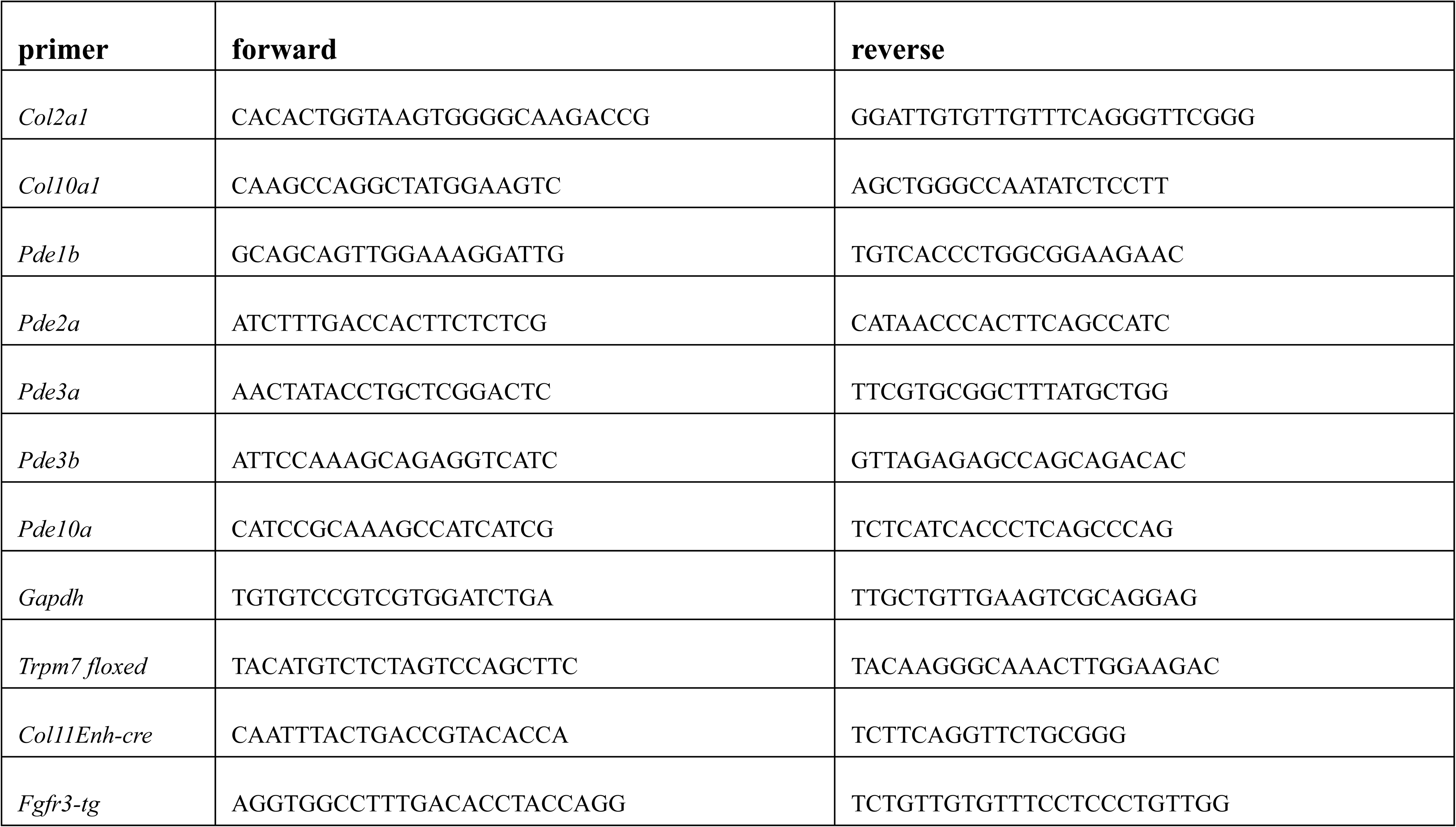
Primers used in this study.

## REFERENCES

Ahrens MJ, Dudley AT (2011) Chemical Pretreatment of Growth Plate Cartilage Increases Immunofluorescence Sensitivity. Journal of Histochemistry & Cytochemistry 59: 408–418

Bartels CF, Bükülmez H, Padayatti P, Rhee DK, van Ravenswaaij-Arts C, Pauli RM, Mundlos S, Chitayat D, Shih L-Y, Al-Gazali LI et al (2004) Mutations in the Transmembrane Natriuretic Peptide Receptor NPR-B Impair Skeletal Growth and Cause Acromesomelic Dysplasia, Type Maroteaux. The American Journal of Human Genetics 75: 27–34

Berendsen AD, Olsen BR (2015) Bone development. Bone 80: 14–18

Calderone V (2002) Large-conductance, ca(2+)-activated k(+) channels: function, pharmacology and drugs. Curr Med Chem 9: 1385–1395

Ceddia RP, Liu D, Shi F, Crowder MK, Mishra S, Kass DA, Collins S (2021) Increased Energy Expenditure and Protection From Diet-Induced Obesity in Mice Lacking the cGMP-Specific Phosphodiesterase PDE9. Diabetes 70: 2823–2836

Dvir L, Srour G, Abu-Ras R, Miller B, Shalev SA, Ben-Yosef T (2010) Autosomal-Recessive Early-Onset Retinitis Pigmentosa Caused by a Mutation in PDE6G, the Gene Encoding the Gamma Subunit of Rod cGMP Phosphodiesterase. The American Journal of Human Genetics 87: 258–264

Fleig A, Chubanov V (2014) Trpm7. In: Mammalian Transient Receptor Potential (TRP) Cation Channels, pp. 521–546.

Heckman PRA, Blokland A, Bollen EPP, Prickaerts J (2018) Phosphodiesterase inhibition and modulation of corticostriatal and hippocampal circuits: Clinical overview and translational considerations. Neuroscience & Biobehavioral Reviews 87: 233–254

Ichimura A, Miyazaki Y, Nagatomo H, Kawabe T, Nakajima N, Kim GE, Tomizawa M, Okamoto N, Komazaki S, Kakizawa S et al (2023) Atypical cell death and insufficient matrix organization in long-bone growth plates from Tric-b-knockout mice. Cell Death Dis 14: 848

Kake T, Kitamura H, Adachi Y, Yoshioka T, Watanabe T, Matsushita H, Fujii T, Kondo E, Tachibe T, Kawase Y et al (2009) Chronically elevated plasma C-type natriuretic peptide level stimulates skeletal growth in transgenic mice. Am J Physiol Endocrinol Metab 297: E1339–1348

Khan S, Basit S, Khan MA, Muhammad N, Ahmad W (2016) Genetics of human isolated acromesomelic dysplasia. European Journal of Medical Genetics 59: 198–203

Kherallah RY, Khawaja M, Olson M, Angiolillo D, Birnbaum Y (2021) Cilostazol: a Review of Basic Mechanisms and Clinical Uses. Cardiovascular Drugs and Therapy 36: 777–792

Kyle BD, Braun AP (2014) The regulation of BK channel activity by pre- and post-translational modifications. Frontiers in Physiology 5

Legeai-Mallet L (2016) C-Type Natriuretic Peptide Analog as Therapy for Achondroplasia. In: Advanced Therapies in Pediatric Endocrinology and Diabetology, pp. 98–105.

Mackie EJ, Ahmed YA, Tatarczuch L, Chen KS, Mirams M (2008) Endochondral ossification: How cartilage is converted into bone in the developing skeleton. The International Journal of Biochemistry & Cell Biology 40: 46–62

Miyazaki Y, Ichimura A, Kitayama R, Okamoto N, Yasue T, Liu F, Kawabe T, Nagatomo H, Ueda Y, Yamauchi I et al (2022) C-type natriuretic peptide facilitates autonomic Ca2+ entry in growth plate chondrocytes for stimulating bone growth. eLife 11

Mouser VH, Melchels FP, Visser J, Dhert WJ, Gawlitta D, Malda J (2016) Yield stress determines bioprintability of hydrogels based on gelatin-methacryloyl and gellan gum for cartilage bioprinting. Biofabrication 8: 035003

Nakao K, Arai H, Komatsu Y, Kanamoto N, Miura M, Murao N, Kondo E, Fujii T, Kitamura H, Yasoda A (2009) Systemic Administration of C-Type Natriuretic Peptide as a Novel Therapeutic Strategy for Skeletal Dysplasias. Endocrinology 150: 3138–3144

Naski MC, Colvin JS, Coffin JD, Ornitz DM (1998) Repression of hedgehog signaling and BMP4 expression in growth plate cartilage by fibroblast growth factor receptor 3. Development 125: 4977–4988

Omori K, Kotera J (2007) Overview of PDEs and Their Regulation. Circulation Research 100: 309–327

Qian N, Ichimura A, Takei D, Sakaguchi R, Kitani A, Nagaoka R, Tomizawa M, Miyazaki Y, Miyachi H, Numata T et al (2019) TRPM7 channels mediate spontaneous Ca(2+) fluctuations in growth plate chondrocytes that promote bone development. Sci Signal 12

Robinson JW, Dickey DM, Miura K, Michigami T, Ozono K, Potter LR (2013) A human skeletal overgrowth mutation increases maximal velocity and blocks desensitization of guanylyl cyclase-B. Bone 56: 375–382

Sadek MS, Cachorro E, El-Armouche A, Kämmerer S (2020) Therapeutic Implications for PDE2 and cGMP/cAMP Mediated Crosstalk in Cardiovascular Diseases. International Journal of Molecular Sciences 21

Samidurai A, Xi L, Das A, Iness AN, Vigneshwar NG, Li P-L, Singla DK, Muniyan S, Batra SK, Kukreja RC (2021) Role of phosphodiesterase 1 in the pathophysiology of diseases and potential therapeutic opportunities. Pharmacology & Therapeutics 226

Savarirayan R, Tofts L, Irving M, Wilcox W, Bacino CA, Hoover-Fong J, Ullot Font R, Harmatz P, Rutsch F, Bober MB et al (2020) Once-daily, subcutaneous vosoritide therapy in children with achondroplasia: a randomised, double-blind, phase 3, placebo-controlled, multicentre trial. The Lancet 396: 684–692

Siuciak JA, McCarthy SA, Chapin DS, Martin AN, Harms JF, Schmidt CJ (2008) Behavioral characterization of mice deficient in the phosphodiesterase-10A (PDE10A) enzyme on a C57/Bl6N congenic background. Neuropharmacology 54: 417–427

Staines KA, Brown G, Farquharson C (2019) The Ex Vivo Organ Culture of Bone. Methods Mol Biol 1914: 199–215

Stratakis CA, Manganiello V, Ahmad F, de Alexandre RB, Levy I, Horvath A, Bimpaki E, Faucz FR, Azevedo MF (2014) Clinical and Molecular Genetics of the Phosphodiesterases (PDEs). Endocrine Reviews 35: 195–233

Suga S, Nakao K, Hosoda K, Mukoyama M, Ogawa Y, Shirakami G, Arai H, Saito Y, Kambayashi Y, Inouye K et al (1992) Receptor selectivity of natriuretic peptide family, atrial natriuretic peptide, brain natriuretic peptide, and C-type natriuretic peptide. Endocrinology 130: 229–239

Teixeira CC, Agoston H, Beier F (2008) Nitric oxide, C-type natriuretic peptide and cGMP as regulators of endochondral ossification. Developmental Biology 319: 171–178

Tsang SH, Gouras P, Yamashita CK, Kjeldbye H, Fisher J, Farber DB, Goff SP (1996) Retinal degeneration in mice lacking the gamma subunit of the rod cGMP phosphodiesterase. Science 272: 1026–1029

Ueda Y, Yasoda A, Hirota K, Yamauchi I, Yamashita T, Kanai Y, Sakane Y, Fujii T, Inagaki N (2019) Exogenous C-type natriuretic peptide therapy for impaired skeletal growth in a murine model of glucocorticoid treatment. Sci Rep 9: 8547

Wang Y, Spatz MK, Kannan K, Hayk H, Avivi A, Gorivodsky M, Pines M, Yayon A, Lonai P, Givol D (1999) A mouse model for achondroplasia produced by targeting fibroblast growth factor receptor 3. Proc Natl Acad Sci U S A 96: 4455–4460

Yamazaki D, Tabara Y, Kita S, Hanada H, Komazaki S, Naitou D, Mishima A, Nishi M, Yamamura H, Yamamoto S et al (2011) TRIC-A Channels in Vascular Smooth Muscle Contribute to Blood Pressure Maintenance. Cell Metabolism 14: 231–241

Yasoda A, Komatsu Y, Chusho H, Miyazawa T, Ozasa A, Miura M, Kurihara T, Rogi T, Tanaka S, Suda M et al (2004) Overexpression of CNP in chondrocytes rescues achondroplasia through a MAPK-dependent pathway. Nat Med 10: 80–86

Young RA, Ward A (1988) Milrinone. A preliminary review of its pharmacological properties and therapeutic use. Drugs 36: 158–192

Phenotype of PDE5a null mice (2011) International Mouse Phenotyping Consortium MGI:2651499 (https://www.mousephenotype.org/data/genes/MGI:2651499) [DATASET]

Phenotype of PDE3b null mice (2011) International Mouse Phenotyping Consortium MGI:1333863 (https://www.mousephenotype.org/data/genes/MGI:1333863) [DATASET]

